# Prophage and SaPI diversity in local clinical *Staphylococcus aureus* reveal hidden drivers of virulence

**DOI:** 10.1101/2025.08.06.669032

**Authors:** Alan Aguayo-González, Irma Martínez-Flores, Patricia Bustos, Rosa I Santamaría, Roberto Cabrera-Contreras, Areli Martínez, Rodrigo Ibarra-Chávez, Víctor González

**Affiliations:** Centro de Ciencias Genómicas, Universidad Nacional Autónoma de México, Cuernavaca, Morelos, México; Laboratorio de Patogenicidad Bacteriana, Departamento de Salud Pública, Facultad de Medicina, Universidad Nacional Autónoma de México, Ciudad de México, México; Instituto Nacional de Ciencias Médicas y Nutrición Salvador Zubirán. Ciudad de México, México; Section of Microbiology, Department of Biology, University of Copenhagen, Denmark; Center for Evolutionary Hologenomics, Globe Institute, University of Copenhagen, Copenhagen, Denmark

## Abstract

Prophages are key drivers of bacterial evolution, genomic variability, virulence, and ecological adaptation. Here, we examined the prophage diversity of 109 *Staphylococcus aureus* clinical isolates from four tertiary care hospitals in Mexico City, integrating comparative analyses with 993 international genomes. Prophages were detected in 97% of the local strains, mirroring the near-ubiquitous presence (99%) observed globally. We identified 216 genomic regions encoding putative prophage functional elements. Using Mitomycin C induction, we recovered 17 temperate phages, 12 of which were functionally active and capable of lysogeny-lysis switching and reinfection. Induced phages lacked virulence or antibiotic resistance genes, in contrast to 55% of the predicted prophages in local *S. aureus* isolates that harbored known virulence factors. Moreover, 19% of the predicted prophages were phage-inducible chromosomal islands (PICIs) related to SaPI1, SaPI2, and SaPIpt1028. These PICIs encode anti-phage defense systems (63%) and virulence genes (27%), such as *tsst-1* and *hlb*. Our findings reveal a complex prophage and PICI landscape in *S. aureus* circulating in Mexican hospitals, with implications for phage– host dynamics, horizontal gene transfer, and the evolution of pathogenic potential across geographically diverse populations.

**Importance:** *Staphylococcus aureus* is a major hospital pathogen whose evolution is driven by mobile genetic elements including prophages and phage-inducible chromosomal islands (PICIs). Although computational analyses predict prophages in nearly all *S. aureus* genomes, we demonstrate that only a small fraction is functionally active. Through experimental induction of clinical isolates from Mexican hospitals, we recovered active prophages that surprisingly lacked virulence genes, in contrast to the predicted prophages that frequently harbor virulence factors. Importantly, we discovered that many predicted “prophages” are PICIs carrying anti-phage defenses and toxins. These findings challenge the current prophage prediction methods and reveal the complex prophage-PICI landscape that shapes horizontal gene transfer and pathogenic evolution in clinical *S. aureus* populations.

## Introduction

*Staphylococcus aureus* is a globally significant opportunistic pathogen responsible for a wide range of infections, from superficial skin abscesses to life-threatening conditions, such as pneumonia, endocarditis, and sepsis. A key feature of *S. aureus* is its remarkable genomic diversity, which is driven mainly by mobile genetic elements (MGEs). These include bacteriophages (phages), phage-inducible chromosomal islands (PICIs), staphylococcal chromosomal cassettes (SCCmec), transposons, insertion sequences, conjugative plasmids, and genomic islands (GIs), all of which contribute to virulence, antibiotic resistance, and adaptation to diverse environments, particularly in hospital settings (1, 2).

Among these MGEs, phages play a crucial role in the horizontal transfer of virulence genes within *S. aureus* populations (1, 3–6). Despite the relatively small number of experimentally characterized *S. aureus* phages, ϕSa3, ϕ11, ϕ80, and ϕMu5 encode key virulence determinants (7–9). These include immune evasion factors (e.g., Sak, Scn, and Chp), Panton-Valentine leukocidin (LukF-PV and LukS-PV), and superantigens (Sea, Sep, and Sec), which are often associated with major epidemic MRSA lineages including clonal complexes CC5, CC8, and CC30 (10–12). However, phages associated with the methicillin-susceptible *S. aureus* (MSSA) lineages remain poorly understood.

Chromosomally integrated prophages are common in *S. aureus*, with most strains harboring one to four such elements (4). Although they rarely encode antibiotic resistance genes, they are predicted to encode virulence-associated genes (6). Furthermore, bioinformatics predictions often identify incomplete and non-inducible prophages, raising questions regarding the fraction of prophages in *S. aureus* that can produce infectious particles.

Experimental studies have shown that treatment of *S. aureus* strains of human and veterinary origin with mitomycin C (MC) can induce prophage activation (13, 14). Notably, these temperate phages, which all belong to the class Caudoviricetes (tailed phages characterized by icosahedral capsids and double-stranded DNA genomes), did not harbor known virulence or resistance genes, highlighting the complexity of phage content in the *S. aureus* genomes.

Importantly, phages also mediate the mobilization of other MGEs, such as PICIs. These, first classified as *S. aureus* pathogenicity islands (SaPIs), encode potent virulence factors. SaPIs can excise and replicate by interacting with a diverse number of phage proteins expressed upon phage infection or induction. SaPI activation leads to capsid assembly and packaging redirection, wherein SaPI DNA is introduced into the phage capsids (2, 15). This mechanism facilitates effective transfer at high frequencies and is recognized as a major route of horizontal gene transfer (16–18).

Given this background, hospital environments can be considered as ecological niches in which *S. aureus* is exposed to intense selective pressures, including antibiotic treatments and host immune responses. Under such conditions, prophages and SaPIs can play a critical role in enhancing bacterial survival and adaptability by benefitting bacterial hosts with the horizontal transfer of genes encoding virulence factors and toxins. However, it remains unclear which prophages circulating in clinical *S. aureus* strains are functionally active and capable of engaging in horizontal gene transfer (HGT) under clinically relevant stress conditions such as antibiotic exposure (19). Thus, we examined the inducibility and mobilization potential of these MGEs to elucidate their impact on the evolution and pathogenesis of hospital-associated *S. aureus*.

In this study, we identified inducible prophages in a collection of *S. aureus* isolated from patients with diverse infection types in tertiary care hospitals in Mexico City. Our results showed that only a subset of the predicted prophages in *S. aureus* genomes were functionally active, that is, capable of producing infectious viral particles and establishing lysogeny in other *S. aureus* strains after MC induction. Consistent with previous reports, these inducible phages did not encode any known virulence or antibiotic resistance genes. However, our analysis also revealed the presence of SaPIs, supporting the notion that phage–SaPI interactions remain a prominent mechanism for the horizontal dissemination of virulence factors within *S. aureus* populations. These findings highlight the complex interplay between phages and other MGEs and underscore the need to further explore their functional roles in clinical contexts.

## Results

### Prophage diversity in *S. aureus* genomes

To explore the extent and structure of the prophage diversity in *S. aureus*, we conducted a systematic analysis of 993 complete genomes retrieved from the NCBI GenBank database as of April 2022 (see Methods and Table S1). These genomes, representing a broad spectrum of globally distributed isolates, were analyzed using the VIBRANT v1.2.1 tool (20), which identifies viral-like regions based on conserved phage signatures, genome architecture, and integration signals. Our analysis revealed that 99% of the *S. aureus* genome (983/993) harbored at least one prophage region, with most strains containing between one and four distinct prophage regions. Only ten genomes lacked detectable prophage elements, underscoring the pervasive nature of prophage integration within the species.

In total, 3279 genomic regions exhibited features consistent with integrated prophages. These regions were subsequently categorized into three quality tiers (high, medium, and low) based on a combination of criteria: genome completeness, presence of hallmark viral genes (such as capsid, tail, DNA polymerase, and terminase proteins), and evidence of site-specific chromosomal integration including attL/attR motifs. This classification allowed us to refine the distinction between putative complete prophages and degraded or fragmented remnants of viral DNA.

Among these, 1972 regions (60%) were classified as either high- or medium-quality prophages, characterized by conserved phage gene modules and genome sizes that closely matched those of experimentally validated *S. aureus* phages available in the NCBI viral genome database (Fig. S1). In contrast, low-quality regions are typically shorter, genetically fragmented, and lack key viral signatures, reflecting either incomplete integration events or ancient prophage decay.

Together, these results revealed a high prevalence and considerable structural variation of prophages across *S. aureus* genomes, showcasing their contribution to the adaptive potential of this species.

### Prophages related to the MRSA isolates

To evaluate whether the prevalence and diversity of prophages differed between methicillin-resistant (*S. aureus*, MRSA) and methicillin-susceptible (*S. aureus*, MSSA) strains, we focused our analysis on high- and medium-quality prophages, excluding fragmented or low-confidence regions (Fig. S1). A statistically significant difference in prophage abundance was observed between MRSA (n = 467) and MSSA genomes (n = 526), with a higher median number of prophages in MRSA strains (*Wilcoxon test*, p = 3.6 × 10⁻¹⁷; Fig. 1).

**Figure 1.**
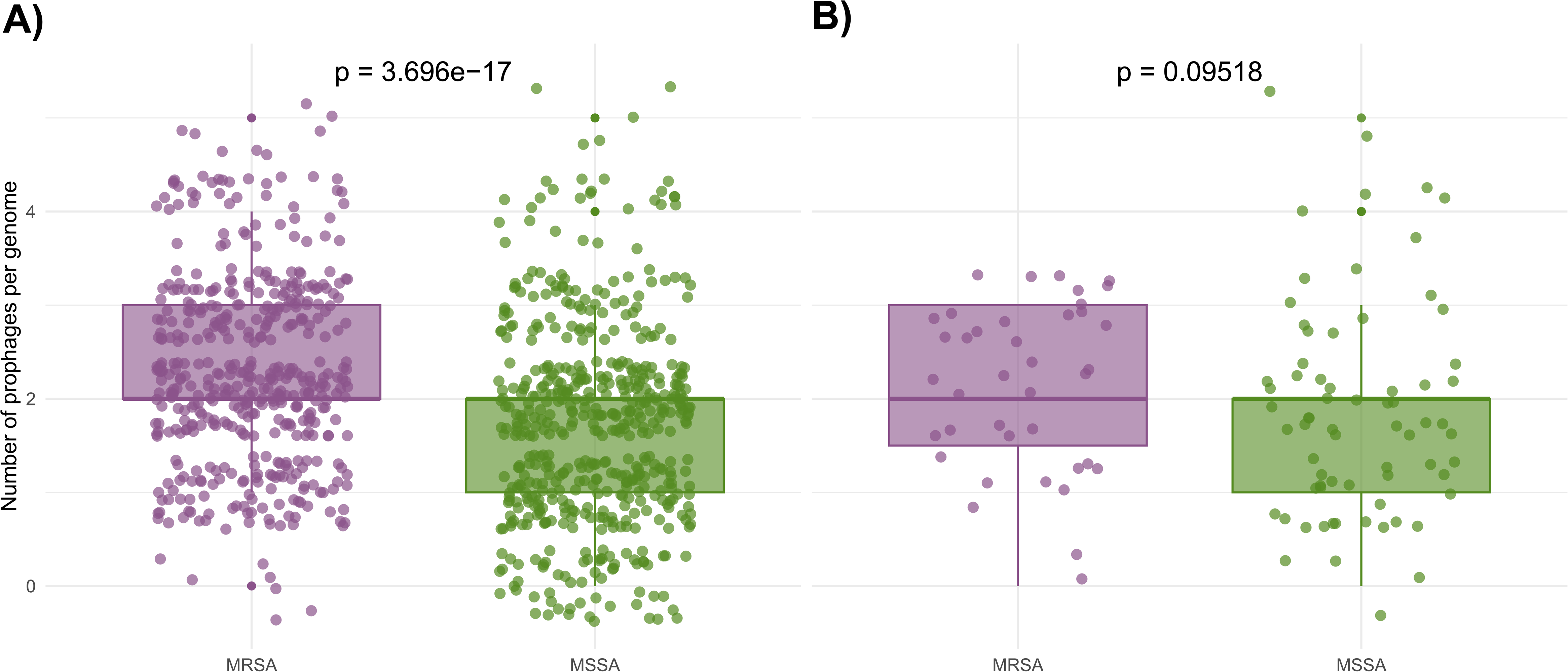
Distribution of prophage content in methicillin-resistant and-susceptible *S. aureus* isolates. Box plots show the number of prophages per genome in MRSA (purple) and MSSA (green) strains from international A) and local collections B). High- and medium-quality VIBRANT predictions were included (see Methods section). The Wilcoxon tests for pairs of data comparisons are shown for each comparison.

These differences suggest a potential relationship between prophage abundance and methicillin resistance but do not necessarily reflect prophage diversity. To further explore this aspect, we assessed prophage diversity by clustering high- and medium-quality prophage sequences using vConTACT2 (21), which groups viral genomes based on their shared protein content and overall similarity.

Prophage sequences were distributed across 124 viral clusters (VCs). Among these, 35 VCs exclusively contained prophages originating from MRSA genomes, 43 contained only prophages from MSSA genomes, and 46 contained prophages from both the backgrounds. Although this distribution suggests that certain prophage lineages are more prevalent in either MRSA or MSSA strains, such patterns should be interpreted with caution. These associations may be driven by clonal population structure, sampling biases, or lineage-specific phage exposure, rather than a direct relationship with the resistance phenotype.

### Local prophage diversity in clinical *S. aureus* isolates

To investigate prophage diversity within a regional context, we analyzed 109 genomes of *S. aureus* clinical isolates obtained from patients with infectious diseases treated in hospitals across Mexico City (Table S2). This dataset, downloaded from GenBank, provides a representative overview of the genomic diversity of *S. aureus* strains circulating in local health care settings.

Prophage regions were detected in 97% (106/109) of genomes, underscoring their widespread occurrence. The three genomes without detectable prophages, INPER770, INPER772, and INPER775, corresponded to MSSA isolates belonging to clonal complex 5 (CC5), a lineage also observed to harbor prophages in other isolates, suggesting that the absence may be strain-specific rather than lineage-associated.

A total of 519 prophage regions were identified, including 116 high-quality, 100 medium-quality, and 303 low-quality regions. As in the global genome dataset, high- and medium-quality prophages in local isolates showed genome size distributions comparable to those of experimentally characterized *S. aureus* phages as well as those predicted in international genomes (Fig. S1).

When stratifying local isolates by methicillin resistance status, MRSA genomes appeared to contain slightly more high-quality prophage regions than the MSSA genomes (Fig. 1). However, this difference was modest and should be interpreted cautiously, as it may reflect variability at the strain level rather than a consistent association between the resistance phenotype and prophage content.

### Prophage induction in clinical *S. aureus* isolates

Previous studies have shown that computational tools can reliably predict the presence of multiple phages and phage-related elements in the *S. aureus* genomes (3, 4, 22). However, these *in silico* approaches offer limited insights into the functional state of predicted prophages, particularly their ability to excise and produce infectious phage particles upon induction.

To address this limitation, we experimentally assessed the presence of inducible prophages in 109 clinical *S. aureus* isolates by treating bacterial cultures with mitomycin C (0.5 μg/ml), a DNA-damaging agent known to activate the SOS response, thereby triggering prophage induction (23, 24). Following treatment, 44% of the strains (48/109) exhibited a marked reduction in growth compared with the untreated controls (Fig. 2), consistent with potential prophage induction and associated cellular stress.

**Figure 2.**
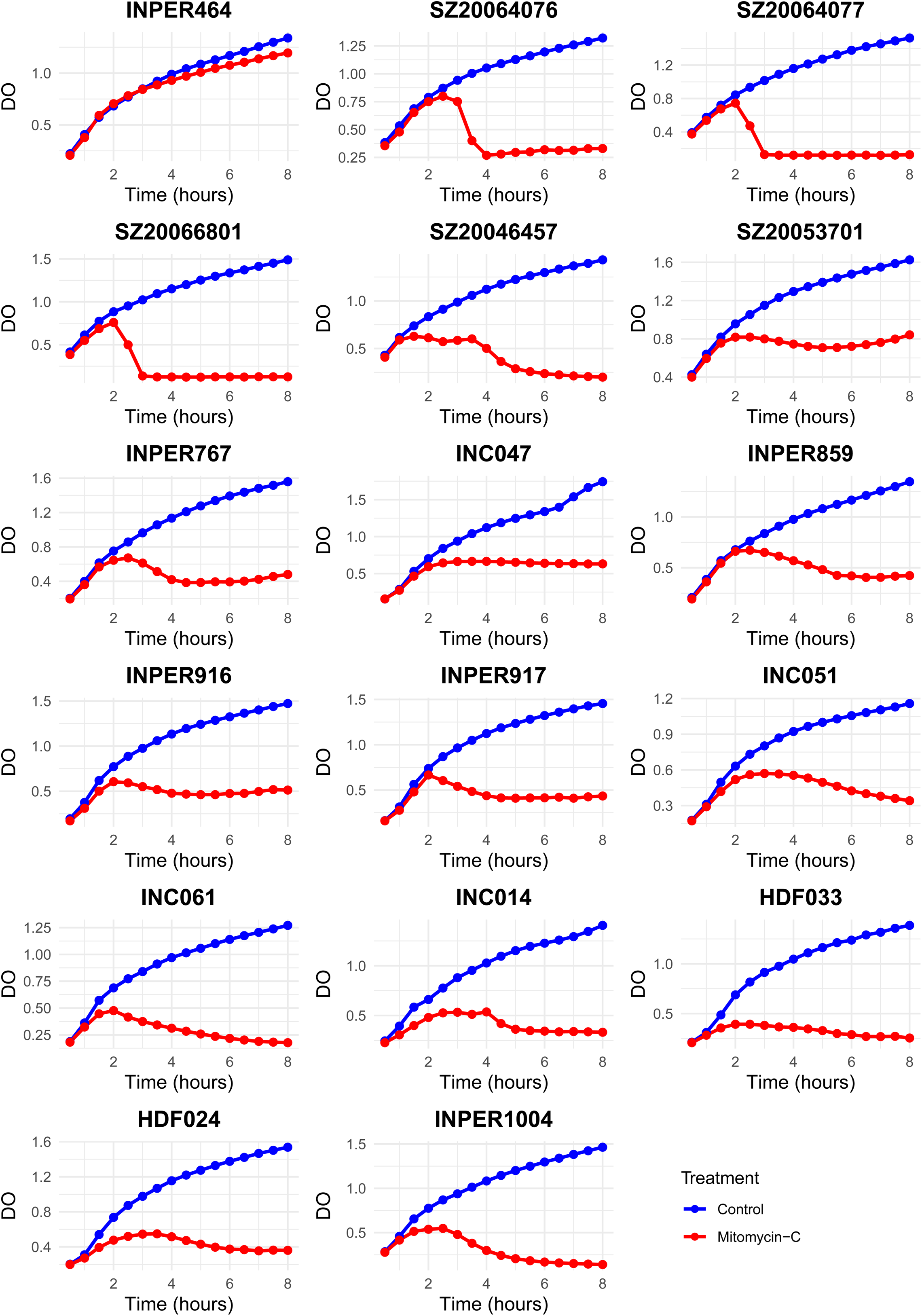
*S. aureus* growth kinetics with and without mitomycin C treatment. *S. aureus* growth in broth cultures was recorded by the change in optical density (DO560) over incubation time (8 hours). Blue curves represent control growth (no treatment), whereas red curves represent growth with mitomycin-C. The first graph (top-left) shows the control strain, INPER464, unaffected by mitomycin-C. The remaining graphs depict the strains in which 17 phages were successfully induced in this study.

As we lacked a universal *S. aureus* indicator strain to directly test for the presence of infectious particles in each case, we employed an indirect approach using spot assays. A subset of 40 *S. aureus* strains was selected as a panel of potential recipients and each of the 48 mitomycin-treated lysates was tested against this panel using double-layer agar spot tests. Notably, 58% of the lysates (28/48) exhibited detectable lytic activity against at least one of the 40 strains, producing either clear or turbid plaques, which is indicative of infectious phage release.

To recover individual phage particles or virions, lysates were subjected to three rounds of plaque purification, resulting in the isolation of 17 distinct phages (Fig. S2). Notably, 11 lysates failed to produce plaques in the spot assay. This suggests that either no phage was present in the lysate or that there was no host strain in the panel that served as a receptor. These findings confirm that a substantial proportion of clinical *S. aureus* strains harbor prophages that are not only genetically identifiable but also functionally inducible, highlighting their potential role in horizontal gene transfer and bacterial lysis under stress conditions.

### Lysogenic properties of the induced prophages

To investigate whether the 17 purified phages could become lysogens in new *S. aureus* host strain and subsequently transitioning back to the lytic cycle, we conducted a series of lysogenizing assays. Briefly, phage lysates were spotted onto lawns of sensitive *S. aureus* strains, and viable colonies emerging from the lysis zones were recovered and tested for phage resistance as an initial indicator of potential lysogen formation (see Methods).

Based on the previously determined host ranges of the 17 phages (Fig. S3), lytic growth in nutrient broth, and spot assays, we selected 31 different phage-host strain pairs for lysogeny testing (17 phages against one to two strains per phage when applicable). From each lysis zone in the spot assay, we isolated up to three independent colonies (denoted L1–L3) per phage-host combination (Fig. 3A) and evaluated their susceptibility to the corresponding phage using spot assays. Among the 31 phage-host combinations tested, in 11 cases, all three colonies exhibited resistance to the original phage, whereas in seven cases, resistance was observed in only one or two of the recovered colonies. In the remaining 13 combinations, no resistant colonies were recovered after phage exposure.

**Figure 3.**
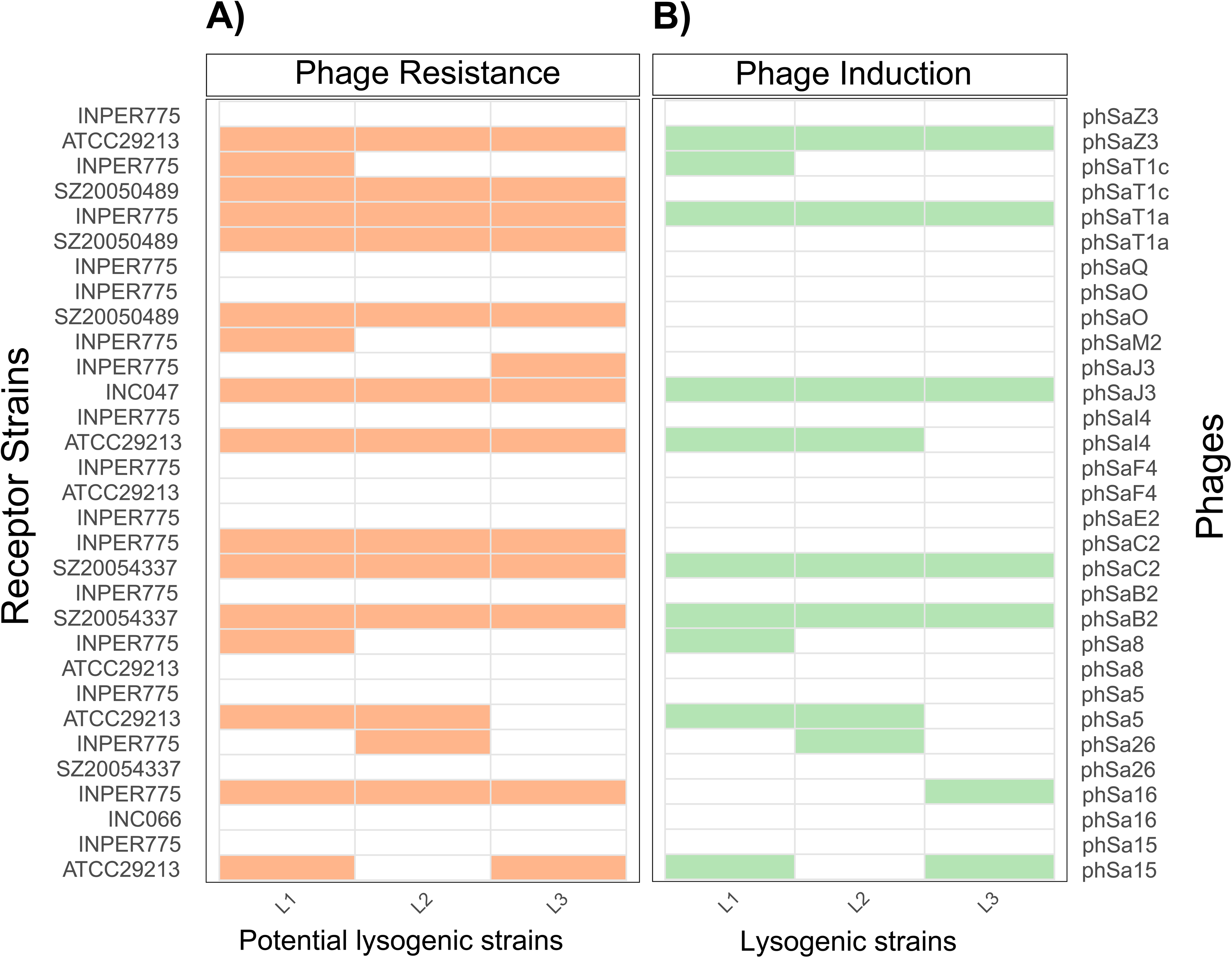
Phenotypes of lysogen strains obtained experimentally. In panels A and B, receptor strains are indicated on the left and phages are indicated on the right. The phenotypes of the three lysogen candidates (L1-to L3, bottom) are indicated at the top of the panel. A) Phage-resistant phenotype: yellow indicates resistance to phage infection and white indicates susceptibility. B) Prophage induction: green indicates successful mitomycin prophage induction and white indicates the absence of prophage induction.

To determine whether phage resistance was due to stable lysogeny rather than alternative mechanisms (e.g., spontaneous mutation or receptor loss), all putatively resistant colonies (n = 42) were treated with mitomycin C, a known prophage inducer. Phage induction was observed in 60% of these colonies (25/42), as evidenced by the production of phage particles that were able of lysing the original non-lysogenic host strain (Fig. 3B). These induced phages were also tested against putative lysogenic strains, confirming that the newly formed lysogens were immune to superinfection by the same phage (Fig. S4B-C), which is consistent with typical lysogenic behavior. Overall, 12 of the 17 phages demonstrated capacity to infect and integrate into new host genomes, establish a stable lysogenic state, re-induce entry into the lytic cycle, and infect other strains. No resistant or lysogenic strains were obtained for phSaE2, phSaF4, phSaM2, phSaO, or phSaQ (Fig. 3A–B). Additionally, in some cases, resistance was observed without evidence of phage induction, suggesting that alternative resistance mechanisms, such as mutations or anti-phage defense mechanisms, may underlie the observed phenotype.

### Core lysogeny genes in induced prophages

To understand the genetic basis of lysogeny in *S. aureus*, we obtained the genomic sequences of 17 induced phages. The genomes had an average size of 44.02 ± 1.53 kb and an average GC content of 34.48% (Table S3). The open reading frames (ORFs) were then predicted and annotated. Across the 17 phage genomes, 1,290 ORFs were identified, with individual genomes encoding between 74 and 81 ORFs (average: 78 ORFs per genome). Approximately 46% of the predicted ORFs were annotated as hypothetical proteins, highlighting a high proportion of phage genes with currently unknown functions. A substantial number of genes have been found to be associated with lysogeny-related functions, highlighting their temperate lifestyle and genetic potential for integration and regulation within host genomes. Notably, 70 ORFs were annotated as transcriptional regulators, indicating a rich repertoire of regulatory elements that may be involved in maintaining lysogeny or controlling the switch to the lytic cycle. Additionally, 19 transcriptional repressors and 16 anti-repressors (including 7 annotated as anti-repressor Ant) were identified, further supporting the presence of systems that modulate phage activation. Core elements of the integration-excision machinery were also prevalent, with 17 integrases and 6 excisionases detected across the genomes. Interestingly, specialized repressors, such as Arc-like (n=2) and CI-like repressors (n=1), were also observed (Table S4), suggesting the conservation of regulatory modules, such as those found in classical temperate phages. Supporting their integrative capacity, the *att* site, a hallmark of site-specific recombination, was identified in 16 of the 17 phages, further confirming their lysogenic nature (Fig. S2, Table S3).

### Prevalence of virulence factors in the prophages of clinical *S. aureus isolates*

To investigate the distribution of virulence-associated genes in prophages identified within clinical *S. aureus* isolates, we analyzed the predicted prophages from 109 local genomes, alongside the 17 experimentally recovered inducible phages, in the context of previously described *Staphylococcus* phages. All prophage elements were grouped based on the shared protein family content using vConTACT2 (21) (see Methods), enabling viral classification and comparative analysis. This approach includes 789 viral elements, comprising 556 known phages from public databases, 216 predicted prophages from local genomes, and 17 inducible phages isolated in this study. This analysis resolved these elements into 38 viral clusters (VCs) (Fig. 4). Fourteen clusters were assigned to recognized ICTV genera, including *Triavirus*, *Phietavirus*, *Dubowvirus*, *Biseptimavirus*, and *Peeveelvirus*. Inducible phages from this study were primarily affiliated with *Triavirus*, *Phietavirus*, and *Dubowvirus*, while *Biseptimavirus* and *Peeveelvirus* were represented exclusively by predicted prophages from the local genomes.

**Figure 4.**
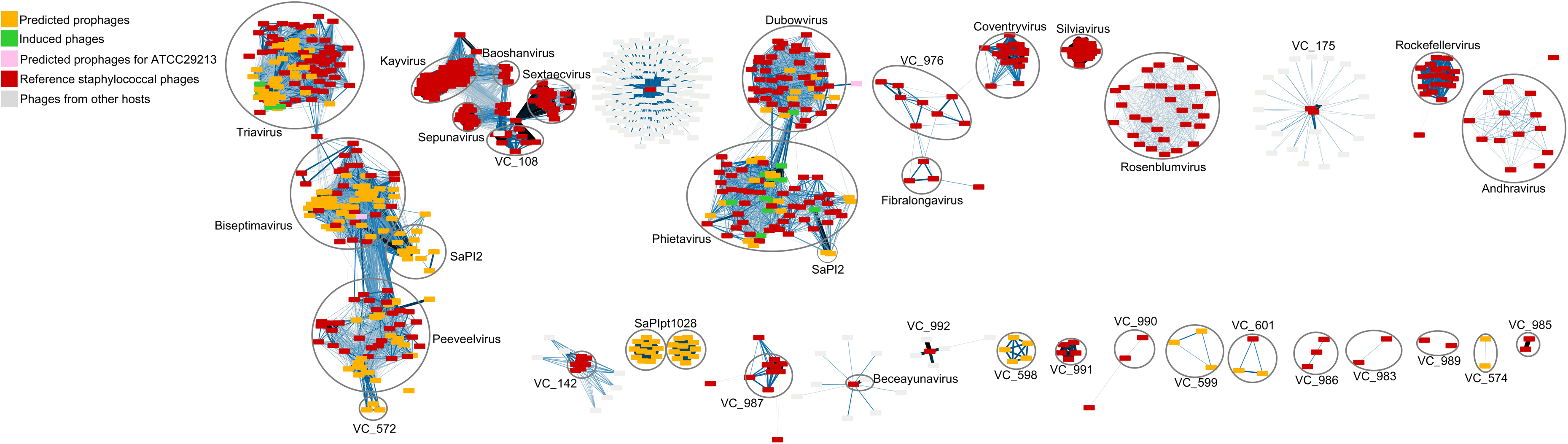
Viral Clusters (VCs) of *S. aureus* phages and prophages. Cytoscape network representation of viral clusters (VCs) based on shared protein family content, as determined by vConTACT2 (21). Each node represents a phage or prophage genome, and the edges indicate similarities in protein content. Known VCs were identified with the names of the VCs and the genus name from the ICTV when available. VCs identified as SaPISs are encircled. Singletons were not included in the network. The predicted prophages from 109 clinical *S. aureus* genomes are shown as orange squares, whereas the 17 inducible prophages isolated in this study are highlighted in green. Reference phages retrieved from the NCBI Viral Genome Database are shown in red (see inset).

To explore the functional potential of these prophages, we screened VCs for virulence-associated genes using Abricate (see Methods section). Several clusters affiliated with *Triavirus*, *Biseptimavirus*, and *Peeveelvirus* encoded known virulence factors. For example, *Triavirus* prophages harbored genes such as elastin-binding protein (Ebp) and the LukS/LukF subunits of Panton-Valentine leukocidin. Members of the *Peeveelvirus* family carry genes encoding staphylokinase (Sak), enterotoxin A (Sea), and beta-hemolysin (Hlb). Similarly, *Biseptimavirus* prophages exhibited diverse virulence profiles, including immune evasion factors such as Sak, staphylococcal complement inhibitor (ScN), chemotaxis inhibitory protein (Chp), and Sea and Hlb toxins (Fig. 5).

**Figure 5.**
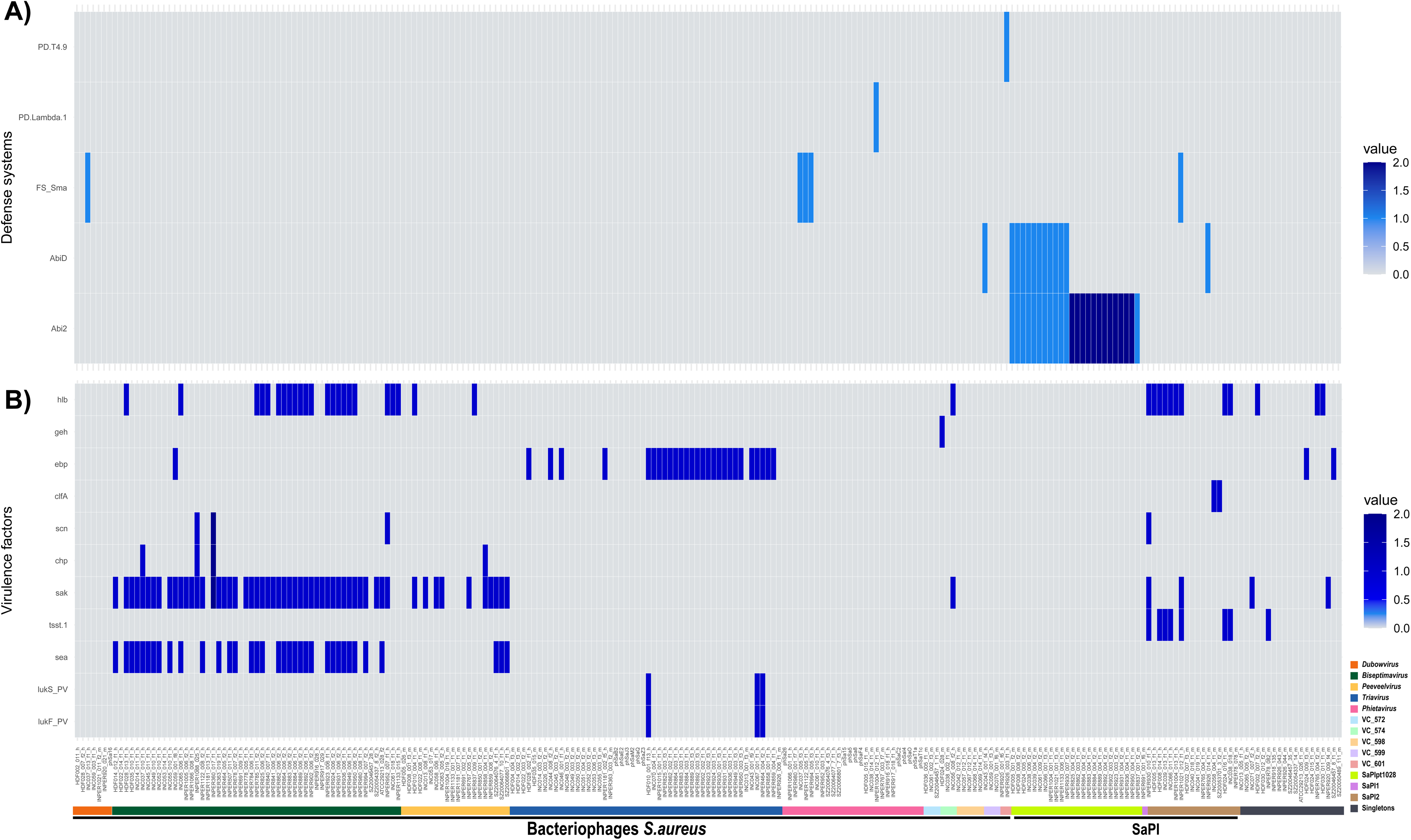
Presence/absence of virulence genes in *S. aureus* phages and prophages. A) Labels on the left indicate anti-phage defense systems, whereas those at the bottom represent phages and prophages. The presence and absence of defense system genes are shown by blue and white rectangles, respectively. Phages were grouped according to their classifications. B) Labels on the left indicate virulence factors, whereas those at the bottom represent phages and prophages. The presence and absence of virulence genes are shown in blue and white, respectively. Phages were grouped according to their classifications.

Interestingly, none of the inducible phages recovered in our experiments were affiliated with *Biseptimaviruses* and *Peeveelviruses,* which were predicted to be widespread genera in the *S. aureus* genomes examined. *Biseptimavirus* was distributed across four subclusters comprising 74 members, whereas *Peeveelvirus* was represented in a single cluster with 39 members. This may reflect the limitation of our experimental induction treatment restrained to a single stressor (MC) or context-specific regulation not assessed under the experimental conditions used here.

In contrast, the inducible prophages associated with *Triavirus*, *Phietavirus*, and *Dubowvirus* generally did not encode identifiable virulence factors, except for some *Triavirus* members (Fig 5). Similarly, phages in the public database ascribed to *Phietavirus* and *Dubowvirus* also lack virulence genes, suggesting a degree of functional conservation within these genera.

Further classification revealed that *Phietavirus* prophages clustered into two distinct subgroups. The first subgroup included 45 members (VC_571_0 to VC_571_3), incorporating nine inducible phages : phSaF4, phSa5, phSa8, phSa15, phSaO, phSaI4, phSaT1a, phSaT1c, and phSaC2. These phages displayed high synteny with the reference phage *Phietavirus pv3MRA*, particularly across structural modules, terminase complexes, and lytic enzymes (Fig. S5C). The second subgroup of *Phietavirus* comprised three subclusters (VC_569_0 to VC_569_2) containing 24 members, including the inducible phage phSa26. This phage exhibited strong genomic similarity to reference phages, such as *Phietavirus ETA*, *StauST3985*, *SAP27*, and *pv80*, with conserved regions involved in DNA processing and capsid formation (Fig. S5D). The genomic divergence between these two *Phietavirus* subgroups, along with limited synteny conservation across non-structural regions, suggests that *Phietavirus* may encompass two distinct genomic lineages, potentially warranting taxonomic re-evaluation within the genus.

### Detection of SaPIs among predicted prophages from clinical isolates

Among the viral clusters predicted through vConTACT2 analysis, we identified two clusters, VC_600 and VC_570, which exhibited no detectable similarity to previously characterized *Staphylococcus* phage sequences. To determine whether these elements represented novel *S. aureus* phages or other MGEs that exploit phage-derived machinery, we conducted a local BLASTn search targeting SaPI-associated integrase genes (Table S5). This analysis revealed that 41 of the predicted prophage regions encoded integrases corresponding to SaPIpt1028, SaPI1, or SaPI2 (Table S6).

Integrases specific to SaPIpt1028 were identified in 24 elements, including representatives from both VC_570 and VC_600 as well as a singleton prophage not assigned to any cluster in the vConTACT2 network (Fig. 4). SaPI1 was identified as a single element (INPER691_001_f6_m), whereas SaPI2-related integrases were detected in 16 predicted elements spanning five viral clusters: VC_1_4, VC_2, VC_3, VC_569, and one singleton. SaPIs showed different virulence patterns according to their classification (Fig. 5). Some SaPI2 elements harbor Hlb (beta hemolysin) and TSS-1 (toxic shock syndrome toxin), which are recognized as being carried by SaPIs and other pathogenicity islands (25, 26). In contrast, islands identified as SaPIpt1028 lacked associated virulence genes, which agrees with previous reports indicating that these islands preferentially carry phage resistance mechanisms (Sma and Abi systems) rather than virulence factors (27) (Table S6). Here, we detected a variant of SaPIpt1028 mainly associated with the Abi2/AbiD systems in two CC lineages, whereas the Sma system was observed to be integrated in other prophages (Fig. 5).

Based on these findings, we reclassified the 41 integrase-containing elements as SaPIs rather than canonical prophages. Consequently, from the initial set of 216 predicted prophage regions, we determined that 175 corresponded to bona fide prophages, whereas 41 represented SaPIs, reflecting the genomic diversity and complexity of phage-related mobile elements in clinical *S. aureus* genomes.

### Phylogenetic relationships and phage/PICI integration patterns in host strains

To understand the relationship between host strains and the presence of different prophage groups, as well as PICIS, a phylogenetic tree was constructed using BPGA v1.3 (28) with data from 109 local strains. The strains were grouped into clades corresponding to their clonal complexes, with CC5, CC8, and CC30 being the most represented. When mapping the various phage groups onto phylogeny, we observed that prophages belonging to the genera *Biseptimavirus* and *Triavirus* were predominantly found in the CC5 and CC8 complexes, especially in methicillin-resistant strains. Regarding pathogenicity islands, SaPIpt1028 showed a broader distribution, being detected in both methicillin-resistant CC8 strains and CC30 MSSA isolates, whereas SaPI2 was exclusively found in CC30 strains (Fig. 6).

**Figure 6.**
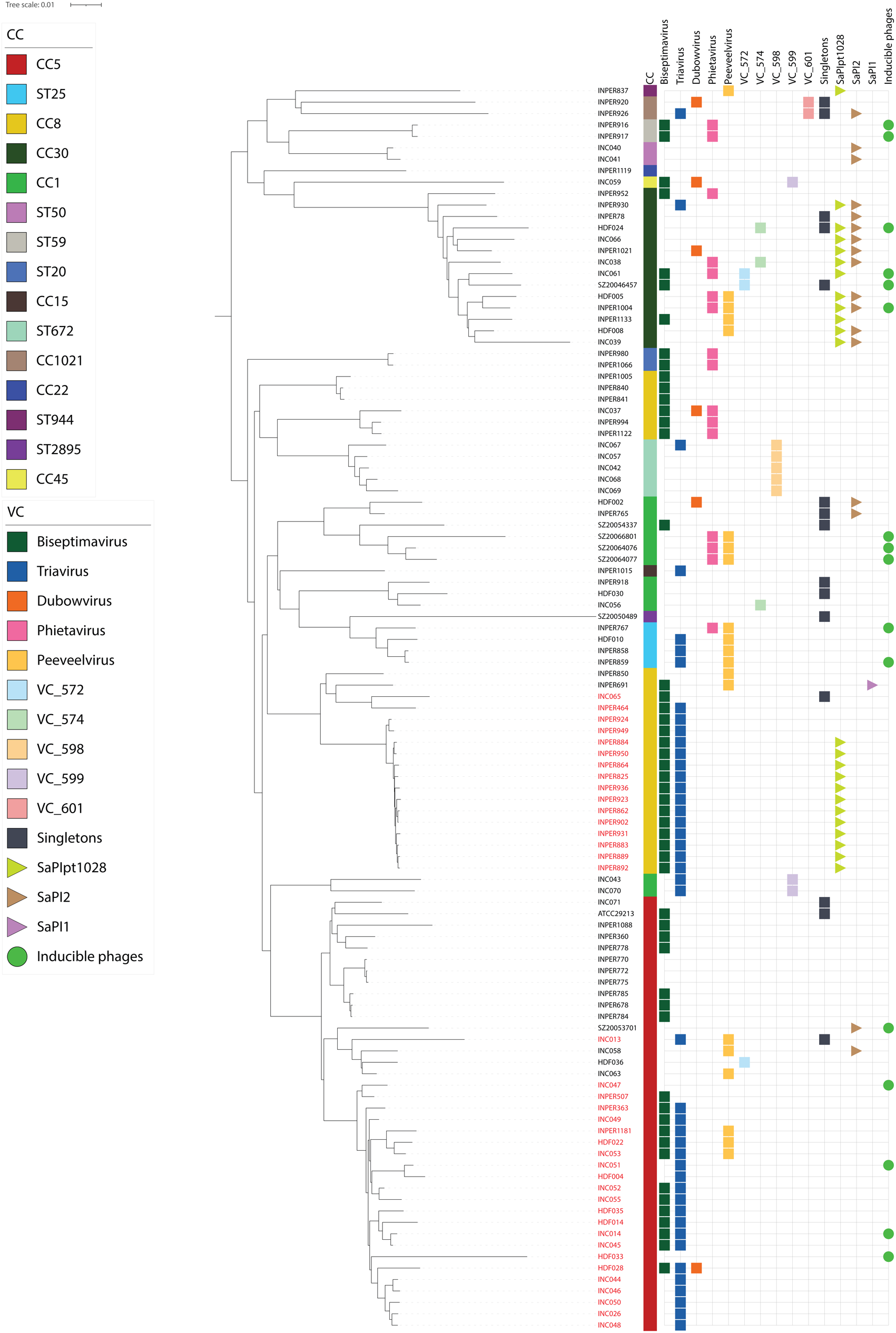
Phylogenetic relationships and prophage integration patterns in host strains. A phylogenetic tree of 109 local *S. aureus* genomes. The tree displays isolate names (black for MSSA and red for MRSA) at the tips, with clonal complexes indicated for each strain. Prophage groups are represented by colored squares corresponding to the predicted prophages found in each strain. Pathogenicity islands are shown as colored triangles and inducible phages are represented by circles. The prophage and pathogenicity island elements were mapped adjacent to their respective host strains in the tree. Color keys for all elements are provided in the figure legends.

## Discussion

This study demonstrated that only a small fraction of the prophages predicted in *S. aureus* genomes was functionally active by MC induction. Although nearly all *S. aureus* strains analyzed carried between one and four prophages, as predicted by the VIBRANT tool, only a limited subset (17 of 109 isolates) produced inducible phages following MC treatment. Remarkably, these induced phages retained the ability to lysogenize new *S. aureus* hosts and could subsequently be reinduced, confirming their functional infectivity and transfer.

Two main explanations could account for the observed low inducibility pattern. First, many prophages may require alternative inducers—chemical, environmental, or stress-related— that differ from MC. For instance, the phage ϕML3 has been shown to respond specifically to pyocyanin, a natural compound produced by *Pseudomonas aeruginosa*, but not to the SOS response caused by MC exposure (29). Second, some prophages may become defective through the loss or mutation of key regulatory or structural genes necessary for excision, replication, or particle formation. Nonetheless, evolutive analyses suggest that prophage genes remain under purifying selection at rates comparable to those of housekeeping genes (30), implying a retained functional potential that might only manifest under specific conditions. These observations highlight the need for a broader search for inducers to complete functional mapping of prophages in *S. aureus*.

The prophages identified in this study did not carry identifiable virulence genes, suggesting a limited direct impact on pathogenicity. This finding aligns with prior reports showing that most *S. aureus* prophages in public repositories (e.g., GenBank and INPHARED) are temperate and lack obvious virulence determinants, as predicted by BACPHLIP. Although the temperate lifestyle of these phages does not exclude their involvement in horizontal gene transfer, *S. aureus* hosts harboring these active prophages often contain virulence and antibiotic resistance genes. This group included four MRSA and thirteen MSSA isolates from patients with confirmed infections, in contrast to the broader genomic pattern in which prophage abundance tends to be higher in MRSA than in MSSA.

The inducible prophages belong to the phage genera *Triavirus*, *Phietavirus*, and *Dubowvirus*, which are generally associated with a low incidence of virulence factor carriage. This contrasts with the genus *Biseptimavirus*, which includes phages known to encode immune evasion genes and toxins. While Panton-Valentine leukocidin (PVL) genes (*lukS-PV* and *lukF-PV*) have been found in some *Triavirus* phages, *Phietavirus* members are not currently known to harbor virulence genes. These patterns reinforce the idea that not all prophages contribute directly to virulence, but may play more subtle roles in shaping the ecology and evolution *of S. aureus*.

Our findings also highlight the limitations of bioinformatic prophage prediction, particularly in draft genome assemblies. For example, a functionally inducible phage was recovered from strain HDF033 despite a lack of high- or medium-quality prophage predictions, likely due to genome fragmentation. Prophage sequence splitting across contigs can hinder accurate prediction and classification, emphasizing the importance of using high-quality, complete genome assemblies for prophage analysis.

An additional layer of complexity arises from the presence of SaPIs, which share structural and functional features with phages and can be mistaken for prophages using standard prediction tools (6). The detection of elements that were initially predicted as prophages but later identified as SaPIs underscores the intricacy of the *S. aureus* mobilome and the need for careful annotation and complementary approaches, such as experimental induction or comparative genomics, to distinguish true prophages from phage-related elements.

Together, our results revealed that only a small proportion of the prophages predicted in *S. aureus* genomes were functionally active under mitomycin C induction. Although most inducible prophages do not carry virulence genes, their persistence and functionality suggest broader roles in host adaptation, competition, and genetic exchange. These findings underscore the importance of combining bioinformatics and experimental approaches to fully understand the dynamics of prophages in clinically relevant *S. aureus* populations and to explore the potential of alternative inducers beyond classical DNA-damaging agents.

## Materials and Methods

### *S. aureus* isolates and global genome dataset

*S. aureus* isolates were obtained from the collection of the Faculty of Medicine of the National Autonomous University of Mexico (FM-UNAM) and from the National Institute of Medical Sciences and Nutrition Salvador Zubirán (INCMNSZ). *S. aureus* isolates (101) from FM-UNAM were previously reported and the complete genomes were retrieved from GenBank (Table S2). The genomes of the *S. aureus* isolates from INCMNSZ were *de novo* sequenced (see below).

For global prophage diversity analysis, 993 complete *S. aureus* genomes were retrieved from the NCBI GenBank database as of April 2022. Genomes were selected based on the assembly quality criteria, including complete genomes, to ensure high-quality sequence data for accurate prophage identification and characterization. Detailed information on the selected genomes is provided in Table S1. MRSA and MSSA classifications were performed using the SCCmec (31) typing tool camlhmp version 1.1.0 (32) with SCCmec_targets schema version 1.2.0 and sccmec_regions schema version 1.2.0.

### Growth conditions for the bacteria, phages, and prophage induction

*Staphylococcus aureus* strains were grown in Lysogenic Broth (LB) medium supplemented with 7 mM CaCl₂ and 10 mM MgSO₄, at 37°C with constant shaking at 210 rpm. Bacto agar was added to the medium when solid plates were required.

Phages were induced by treatment with mitomycin C (0.5 μg/ml) applied during the early logarithmic phase of the cultures. After eight hours of incubation, the cultures were centrifuged at 3000 × g for 10 min and the supernatant was collected and stored at 4°C until further use. Phage detection was performed using a double-layer plaque assay technique with strains from the collection. A total of 200 μl of each sample and 100 μl of overnight culture were added to 4 ml of top agar (soft medium with 0.4% agar melted at 42°C) and poured onto LB plates. The appearance of lytic plaques was visually evaluated, and isolated plaques were recovered and purified in three successive replicates.

### Genomic sequencing of bacteria and bacteriophages

In this study, we sequenced the genomes of eight *S. aureus* isolates from INCMNSZ and 17 phages. Bacterial DNA was obtained using the GenElute Bacterial Genomic DNA Kit (Sigma Aldrich) following standard protocols, with lysostaphin treatment (200 units/ml) for cell lysis and RNase A for RNA removal. DNA quality and concentration were assessed by agarose gel electrophoresis, spectrophotometry (NanoDrop 2000; Thermo Fisher, Waltham, MA, USA), and fluorometry (Qubit dsDNA assay; Thermo Fisher). Whole genome sequencing was performed in BGI Tech Solution using DNBSEQ technology with paired-end 150 bp reads from 350 bp insert libraries, generating 2 Gb per sample. Raw sequencing data underwent quality control and trimming using FastQC v0.11.8 and Trim Galore v0.6.4, followed by *de novo* assembly using SPAdes genome assembler v3.13.1 (33).

Phage DNA was purified following the protocol from the DNA isolation kit for cells and tissues (Roche Life Sciences, Basel, Switzerland), with modifications. The raw sequencing data were subjected to quality control using FastQC v0.11.9 and cleaned using Trim Galore v0.6.7. *De novo* assembly was performed using SPAdes Genome Assembler 3.13.1 (33), MEGAHIT v1.2.9 (34) and Unicycler v0.5.1 (35). Assembly quality metrics were evaluated using QUAST v5.3.0 (36) and the best assembly for each phage was selected based on these metrics. Subsequently, genome completeness was assessed using CheckV 1.0.1 (37).

### Prophage prediction

The prophage content prediction in S. aureus genomes was conducted using the bioinformatics tool VIBRANT (Virus Identification By iteRative ANnoTation) v1.2.1 (20). Default parameters were applied, considering prophage sequences with a minimum length of 1000 bp and an open reading frame (ORF) count of four or more.

### Genome Annotation and Viral Cluster Construction

The prediction and annotation of open reading frames (ORFs) for the isolated phages and predicted prophages were performed using Pharokka v1.4.0 (38), followed by manual curation. Attachment sites were identified using PHASTER (39).

To construct the phage similarity network, viral clusters (VCs) were generated using vConTACT v.2 (21). This network included 789 isolated phages and predicted prophages for *Staphylococcus*, of which 556 were previously reported phages from public databases (NCBI/ICTV/INPHARED), 216 were predicted prophages from local genomes, and 17 were experimentally recovered phages from this study. vConTACT2 groups similar proteins into protein clusters (PCs) using the Markov Clustering Algorithm (MCL) and calculates the maximum likelihood of shared PCs (edges) between genomes (nodes), resulting in a bipartite network. VCs were identified using Cluster ONE with a significance threshold of 60. The resulting network was visualized using Cytoscape v.3.10.2 (40).

### Lysogens isolation

Based on host range analysis, 31 phage-host combinations were selected for lysogeny screening according to specific selection criteria. Combinations were chosen based on their lytic activity during liquid culture growth and the formation of distinct lysis zones in spot assays, but where turbidity (resistant cells) began to appear after repeated passages or extended incubation times. Phage lysates were spotted (10 μl) onto the lawns of the sensitive strains. Following 12 h of incubation at 37°C, viable bacterial cells were recovered from the central region of the lysis zones using sterile bacteriological loops and subsequently streaked for isolation. Three independent colonies (designated L1–L3) were collected and purified through three successive passages on solid medium.

Isolated colonies were cultured in liquid medium and subjected to spot tests to assess their resistance to infection (immunity to superinfection) using the parental phage from which they were derived. Putative lysogenic isolates were further evaluated for prophage induction capacity through treatment with mitomycin C (0.5 μg/ml) for eight hours to assess the potential of lysogenic phages to enter the lytic cycle.

### Identification of SaPIS

Pathogenicity islands (SaPIS) were searched for in the high- and medium-quality prophage predictions obtained using VIBRANT. The complete genomic sequences of all 216 predicted prophages were queried using BLASTp against a curated database of the SaPI integrase genes (Table S5). The BLASTp parameters were set to a minimum of 95% amino acid identity and coverage alignment. The predicted prophages that met these criteria were redefined as SaPIs.

### Phylogenetic analysis and mobile element mapping

Phylogenetic analysis of 109 local S. aureus strains was performed using BPGA v1.3 (28) with the core genome approach, identifying 2,108 core proteins through comparative genomic analysis. Maximum likelihood phylogenetic reconstruction was conducted using IQ-TREE v2.1.2, with default parameters, implementing the JTT+F+R3 evolutionary model, and 1,000 bootstrap replicates to assess branch support.

Multilocus sequence typing (MLST) was employed for clonal complex (CC) assignment. Sequence types (STs) were primarily determined using the BV-BRC Genomic Annotation Service by comparison with established ST alleles in PubMLST (https://pubmlst.org/). For novel alleles absent from PubMLST, the Center for Genomic Epidemiology MLST service (https://cge.food.dtu.dk/services/MLST/) was applied using default parameters and BV-BRC-derived FASTA files to determine the allelic profiles and corresponding STs. Related STs were subsequently grouped into CCs using the eBURST algorithm (https://pubmlst.org/organisms/staphylococcus-aureus), which clusters strains that share similar allelic profiles.

The resulting phylogenetic trees were drawn and edited using the Interactive Tree of Life program (iTOL v7.2). Prophage groups, SaPIs, and other mobile elements were mapped onto a phylogenetic tree using iTOL annotation features.

### Identification of virulence factors, antibiotic resistance genes, and anti-phage systems

Identification of virulence factors and antibiotic resistance genes was carried out using ABRicate 1.0.1 (41) with default parameters (80% identity and 80% coverage), utilizing the VFDB (Virulence Factor Database) (42) and CARD (Comprehensive Antibiotic Resistance Database)(43), respectively. The search for anti-phage defense systems was performed using DefenseFinder 1.3.0 (44), with default parameters.

### Declaration of use of artificial intelligence tools

During the preparation of this work, the authors used ChatGPT (OpenAI, GPT-4) and PaperPal (Cactus Communications Services Pte, LTD) to correct the English grammar and edit the submitted version. The authors declare that the scientific content is original, and after using these tools, they reviewed and edited the content as needed and took full responsibility for the content of the publication.

## Data availability

All relevant data supporting the findings of this study are available in the main text and supplementary files. Genomic sequences from bacterial strains and phages are publicly available through the National Center for Biotechnology Information (NCBI) with accession numbers detailed in the supplementary material.

## Acknowledgments

This work was supported by the Programa de Apoyo a Proyectos de Investigación e Innovación Tecnológica (PAPIIT-UNAM) IN214019 to R.C. and V. G., and by the research budget of Centro de Ciencias Genómicas, UNAM to VG. R.I.C. was supported by a research grant VIL 588733-Weaponizable satellites from VILLUM FONDEN. A. A-G. is a doctoral student from the Programa de Doctorado en Ciencias Biomédicas, UNAM, with a CONAHCYT fellowship 1023079.

## Supplementary material

**Table S1. General characteristics of 993 complete *S. aureus* genomes from NCBI.**

**Table S2. General characteristics of 109 local *S. aureus* genomes.**

**Table S3. General characteristics of the 17 induced phages in this study. Table S4. Functional annotation of induced phages by Pharokka.**

**Table S5. Reference SaPI integrase protein sequences.**

**Table S6. Results of a BLAST search targeting integrases characteristic of S. aureus pathogenicity islands (SaPIs).**

**Fig. S1. Comparison of predicted prophages with phages reported in NCBI.** A) Genome size comparison of predicted prophages in local strains. B) Genome size comparison of predicted prophages in international strains. High-quality prophages are in red, medium-quality in blue, low-quality in green, and reported phages in black.

**Fig. S2. Functional annotation and genomic organization of 17 inducible *Staphylococcus aureus* prophages.** Genomic maps showing functional classification of predicted ORFs across 17 inducible prophages. Genes are color-coded by functional categories: structure and assembly proteins, DNA processing proteins, repressor/regulator proteins, lysozyme and lysins, metabolic genes, transporter proteins, integrases, maturases/terminases, hypothetical and orphan genes, and transposases.

**Fig. S3. Host range analysis of 17 inducible *Staphylococcus aureus* prophages.** Heatmap showing the lytic activity of 17 inducible prophages (listed vertically) against 48 *S. aureus* clinical strains (listed horizontally). Blue cells indicate successful lysis, while white cells represent absence of lytic activity. The top bar chart displays the number of prophages capable of lysing each strain, with strains ordered from most susceptible (left) to least susceptible (right) to prophage infection. The right bar chart shows the number of strains lysed by each prophage, ordered from most infective (top) to least infective (bottom).

**Fig. S4. Experimental validation of lysogenic conversion.** A) Isolation and selection of potential lysogenic colonies following phage exposure. Individual colonies displaying lysogenic characteristics were isolated for subsequent phenotypic and functional analysis. B) Functional validation of prophage induction from lysogenic strains. Spot test comparing the lytic activity of the original phage and phages induced from lysogenic candidates (L1, L2, L3) against the original non-lysogenic host strain, confirming successful prophage induction and production of viable phage particles. C) Superinfection immunity assay demonstrating prophage-mediated protection. Spot test showing the resistance of lysogenic strain to infection by the original phage and phages induced from the three lysogenic strains (L1, L2, L3). The absence of lysis indicates successful establishment of immunity, confirming that the integrated prophage provides protection against superinfection by homologous phages.

**Fig. S5. Synteny of *S. aureus* bacteriophages.** A) Genome comparison of predicted prophages, induced phages (phSa16), and reported phages from the *Dubowvirus* genus. B) Genome comparison of predicted prophages, induced phages (phSaB2, phSaE2, phSaM2, phSaQ, phSaZ3, and phSaJ3), and reported phages from the *Triavirus* genus. C) Genome comparison of predicted prophages, induced phages (phSaF4, phSa5, phSa8, phSa15, phSaO, phSaI4, phSaT1a, phSaT1c, and phSaC2), and reported phages from the *Phietavirus* genus subgroup 1. D) Genome comparison of predicted prophages, induced phages (phSa26), and reported phages from the *Phietavirus* genus subgroup 2. Functional annotation is represented by a color key, and the percentage of ORF similarity is shown in grayscale.

